# CSDE1 promotes passenger strand cleavage of miR-486

**DOI:** 10.1101/2025.09.09.675073

**Authors:** Arezou Arvand, Yunkoo Ko, Tony Chen, Emily Tsao, Mahammed Zaid Patel, Lucas Hiscock, Louis-Mathieu Harvey, Martin Simard, Kristin Hope, Chanseok Shin, Pavan Kumar Kakumani

**Affiliations:** Department of Biochemistry, Memorial University of Newfoundland, 45 Arctic Ave., St. John’s, NL Canada A1C 5S7; Department of Agricultural Biotechnology, Seoul National University, Seoul 08826, Republic of Korea; Department of Medical Biophysics, University of Toronto, Toronto, Canada; Princess Margaret Cancer Center, University Health Network, Toronto, Canada; Oncology Division, Centre Hospitalier Universitaire de Québec-Université Laval Research Center (L’Hôtel-Dieu de Québec), Quebec City, Québec G1R 3S3, Canada; Laval University Cancer Research Centre, Québec City, Québec G1R 3S3, Canada; Research Center for Plant Plasticity, Seoul National University, Seoul 08826, Republic of Korea; Research Institute of Agriculture and Life Sciences, and Plant Genomics and Breeding Institute, Seoul National University, Seoul 08826, Republic of Korea

**Author notes:** These authors contributed equally (Arezou Arvand, Yunkoo Ko).

## Abstract

Strand separation of the RNA duplex is a critical step in miRNA biogenesis, in which only the guide strand is retained to form the mature miRISC. The erythroid miRNA miR-486-5p uniquely requires AGO2 catalytic slicing of its passenger strand miR-486-3p, yet the factors regulating this process remain poorly understood. Here, we identify the RNA-binding protein CSDE1 as a key cofactor that promotes AGO2-dependent passenger-strand removal and miR-486-5p maturation. CSDE1 interacts with the N-terminal domain of AGO2, essential for strand release. Loss of CSDE1 increased the miR-486-3p/5p ratio, reduced cleavage efficiency, and derepressed miR-486 targets such as FOXO1 and PTEN. Restoring full-length CSDE1 expression restored AGO2-dependent passenger strand cleavage and this function requires the N-terminal cold-shock domain CSD1, which mediates CSDE1 interaction with AGO2. Together, these results demonstrate CSDE1 as a critical regulator of AGO2-mediated strand separation, potentiating miR-486-5p function in target gene silencing associated with in hematological malignancies.

## INTRODUCTION

MicroRNAs (miRNAs) are ∼22 nucleotide non-coding RNAs that regulate gene expression by repressing translation or promoting mRNA decay through base-pairing interactions within the 3′ untranslated regions (UTRs) of target transcripts (Hynes and Kakumani 2024). Canonical miRNA biogenesis begins with the transcription of primary miRNAs (pri-miRNAs), which are processed by Drosha into precursor miRNAs (pre-miRNAs). These are then cleaved by Dicer to form mature miRNA duplexes, which are loaded onto Argonaute (AGO) proteins to assemble the miRNA-induced silencing complex (miRISC) (O’Brien et al. 2018; Bartel 2018). While this sequential pathway is conserved across most miRNAs, alternative processing routes exist. Mitrons, for example, bypass Drosha cleavage via splicing (Westholm and Lai 2011). More strikingly, miR-451 represents the only known vertebrate miRNA that is completely Dicer-independent. Due to its short hairpin structure, pre-miR-451 is directly loaded into AGO2, where slicer activity cleaves the passenger strand, followed by poly(A)-specific ribonuclease (PARN)-mediated trimming to generate the mature form (Kakumani et al. 2023). Similarly, miR-486 maturation depends on AGO2 catalytic activity, which facilitates removal of the passenger strand (miR-486-3p), enabling functional incorporation of the guide strand (miR-486-5p) into RISC (Jee et al. 2018).

A growing number of studies have demonstrated that misregulation of miRNA expression is a hallmark of cancer, including leukemia (Peng and Croce 2016; Lu et al. 2005; Yendamuri and Calin 2009). Distinct leukemia-specific miRNA expression patterns, such as the upregulation of let-7b/miR-223 and the downregulation of miR-128a/b, can discriminate acute myeloid leukemia (AML) from acute lymphoblastic leukemia (ALL) (Dixon-McIver et al. 2008; Mi et al. 2007; Jongen-Lavrencic et al. 2008). Dysregulated miRNAs influence leukemic processes ranging from proliferation and differentiation to chemotherapy resistance (Gabra and Salmena 2017; Trino et al. 2018). For example, reduced expression of let-7b increases levels of the oncogenic fusion protein AML1-ETO, thereby sustaining proliferation and blocking differentiation in AML (Johnson et al. 2021). Altered miRNA levels arise through copy number changes, epigenetic modifications, or disrupted processing steps, with the latter representing an underexplored but important mechanism of subverting miRNA function in cancer (Li et al. 2008, 2013; Senyuk et al. 2013; Jiang et al. 2016). Despite the critical role of miRNAs in leukemogenesis, it remains poorly understood whether aberrant maturation of non-canonical miRNAs contributes to AML pathology.

Among hematopoietic miRNAs, miR-451 and miR-486 are highly enriched in red blood cells and play essential roles in erythropoiesis. miR-451 is abundant in mature erythrocytes, whereas miR-486-5p is enriched in maturing erythroblasts, where it promotes erythroid differentiation and apoptosis by repressing FOXO1 and PTEN (Jee et al. 2018). Loss of miR-486 impairs red blood cell development in mouse models, and its dysregulation is linked to leukemogenesis. miR-486-5p exerts context-dependent roles: it functions as a tumor suppressor in AML and several solid tumors, but can act as an oncogene in specific contexts such as Down syndrome associated leukemias and chronic myeloid leukemia (CML), where it modulates progenitor growth, survival, and drug response (Shaham et al. 2015; Liu et al. 2019; Wang et al. 2015). Interestingly, its expression increases in imatinib-treated CML, suggesting a tumor-suppressive role in the BCR-ABL pathway (Ninawe et al. 2021). More broadly, miR-486-5p is frequently downregulated in diverse malignancies, including lung, ovarian, gastric, and renal cancers, where its reintroduction suppresses tumor growth by inducing apoptosis and caspase-3 activation (Xu et al. 2022; ElKhouly et al. 2020). Importantly, miR-451 and miR-486 act in a partially redundant manner to sustain erythropoiesis. Double knockout mice for miR-451 and miR-486 phenocopy AGO2 catalytic-dead mice, underscoring that these two miRNAs are the predominant physiological substrates of AGO2 slicer activity in vivo (Huang et al. 2021; Kakumani et al. 2023). These findings firmly link the activity of AGO2 to the biogenesis of erythropoietic miRNAs, suggesting that additional cofactors may be required to facilitate efficient strand cleavage and functional maturation.

Cold shock domain containing E1 (CSDE1), a multifunctional RNA-binding protein of the CSD family, has recently emerged as a key regulator of non-canonical miRNA processing. CSDE1 is best known for its roles in translation, mRNA stability, and splicing (Kakumani et al. 2021, 2020, 2023). More recently, CSDE1 has been identified as a component of the miRNA RISC, where it interacts with AGO2, the catalytic core of the complex, to promote efficient miRNA target silencing (Kakumani et al. 2021, 2020). In erythroid progenitors, CSDE1 facilitates the atypical AGO2-dependent maturation of the non-canonical miRNA miR-451, acting as a scaffold that recruits AGO2 to pre-miR-451 and promotes passenger-strand cleavage (Kakumani et al., 2023). These observations raise the possibility that CSDE1, in addition to its established role in miR-451 biogenesis, may also contribute to the AGO2-dependent maturation of miR-486.

Beyond hematopoiesis, CSDE1 exerts diverse, context-dependent roles in disease. In cancer, it functions as an oncogene in melanoma, colorectal cancer, glioma, breast cancer, and pancreatic ductal adenocarcinoma, where high expression promotes invasion, metastasis, and poor prognosis (Ningyu et al. 2018; Fang 2005; Wurth et al. 2016; Guo et al. 2020). In contrast, CSDE1 inactivation serves as a tumor suppressor in pheochromocytomas/paragangliomas (PCCs/PGLs) and oral squamous cell carcinoma (Pedro et al. 2018; Fishbein et al. 2017; Guo et al. 2020), while its depletion sensitizes epithelial ovarian cancer to platinum-based therapy (Huang et al. 2021; Guo et al. 2020). These findings highlight CSDE1 as both a prognostic biomarker and a therapeutic target across malignancies. In non-malignant disorders, CSDE1 expression is reduced in Diamond Blackfan anemia (DBA), impairing erythroid proliferation and differentiation, while de novo mutations are strongly associated with autism spectrum disorders (ASDs) (Xia et al. 2014; Horos et al. 2012). During embryogenesis, CSDE1 is indispensable, with knockout models exhibiting mid-gestation lethality due to placental and neural defects. In human embryonic stem cells, CSDE1 also prevents premature differentiation by stabilizing transcripts such as FABP7 and VIM (Ju Lee et al. 2017; Saltel et al. 2017; Guo et al. 2020). Collectively, these studies establish CSDE1 as a global regulator of cell fate decisions.

Within the hematopoietic system, CSDE1 is especially relevant for non-canonical miRNAs. Similar to miR-451, miR-486-5p requires AGO2-mediated passenger-strand cleavage for maturation. miR-486-5p is highly enriched in blood, where it promotes erythroid differentiation and apoptosis by repressing FOXO1 and PTEN (Jee et al. 2018). Importantly, CSDE1 expression is diminished in acute myeloid leukemia (AML), correlating with impaired erythroid differentiation and leukemic progression (Gíslason et al. 2024). Although CSDE1 has been established as an oncogene in some cancers and a tumor suppressor in others, its role in leukemia has only recently begun to be explored. Together, these findings underscore the significance of investigating the CSDE1– AGO2–miR-486 axis, which links RNA regulation to erythropoiesis, leukemia, and potential therapeutic strategies in hematological malignancies.

Therefore, in this study, we identify CSDE1 as a critical regulator of miR-486 maturation. CSDE1 interacts with the AGO2 N-terminal domain to promote passenger-strand removal, ensuring efficient miR-486-5p activity. Loss of CSDE1 disrupts this process, elevating the miR-486-3p/5p ratio, restoring the expression of the target genes, and impairing erythroid differentiation, while its restoration rescues miRNA function. These findings establish CSDE1 as an essential cofactor in AGO2-dependent miR-486 processing, directly connecting RNA strand separation to erythroid differentiation and leukemic progression.

## RESULTS

### CSDE1 promotes the expression of miR-486-5p in leukemia

To investigate whether CSDE1 promotes the expression of miR-486-5p in leukemia, we first examined multiple tissue types using flow cytometry to measure the expression of miR-486-5p in comparison with miR-486-3p, miR-1468-5p, miR-504-3p, and miR-145-3p. This analysis is particularly relevant because the function of miR-486-5p depends on the efficient clearance (or cleavage) of its passenger strand, miR-486-3p, from the AGO2-bound duplex (Jee et al. 2018). Thus, the relative abundance of miR-486-3p versus miR-486-5p bound to AGO2 determines the functional output of miR-486-5p (with a lower 3p/5p ratio indicating enhanced function). Our results showed that miR-486-5p is highly expressed relative to its passenger strand, as well as compared to the other canonical miRNAs analyzed, in blood compared to other tissue types (Rishik et al. 2025) (Figure S1). These findings suggest that miR-486-5p has greater functional activity in blood owing to its lower 3p/5p ratio.

Next, we performed microarray profiling to compare CSDE1 expression levels in acute myeloid leukemia (AML) with those in normal hematopoietic cells. CSDE1 expression was significantly reduced in AML samples relative to normal hematopoietic cells (Figure S2). Previously, we demonstrated that CSDE1 interacts with AGO2 in erythroid progenitor cells and promotes the non-canonical biogenesis of the tumor suppressor miR-451(Kakumani et al. 2023). Given that miR-486-5p is preferentially expressed in blood compared to miR-486-3p, that its function depends on AGO2-mediated passenger-strand cleavage, and that CSDE1 expression is diminished in AML, we hypothesized that CSDE1 promotes AGO2-dependent biogenesis of miR-486-5p in leukemia. To test this hypothesis, we measured relative miRNA levels in total RNA and AGO2 immunoprecipitates (IP) from control and CSDE1 knockout (KO) MEL cells, focusing on miR-486-5p/3p ratios and using miR-20a-5p/3p as a canonical control. Loss of CSDE1 led to increased accumulation of miR-486-3p without significantly affecting miR-486-5p levels (Figure 1A). Consequently, the miR-486-3p/5p ratio bound to AGO2 was significantly elevated in CSDE1 KO cells compared to controls (Figure 1B). Consistent with this, western blot and miRNA quantification demonstrated increased expression of known miR-486-5p targets, including FoxO1 and PTEN, in CSDE1 KO cells (Figure 1C). These results indicate that CSDE1 plays a critical role in promoting AGO2-mediated miR-486-5p function.

**Figure 1.**
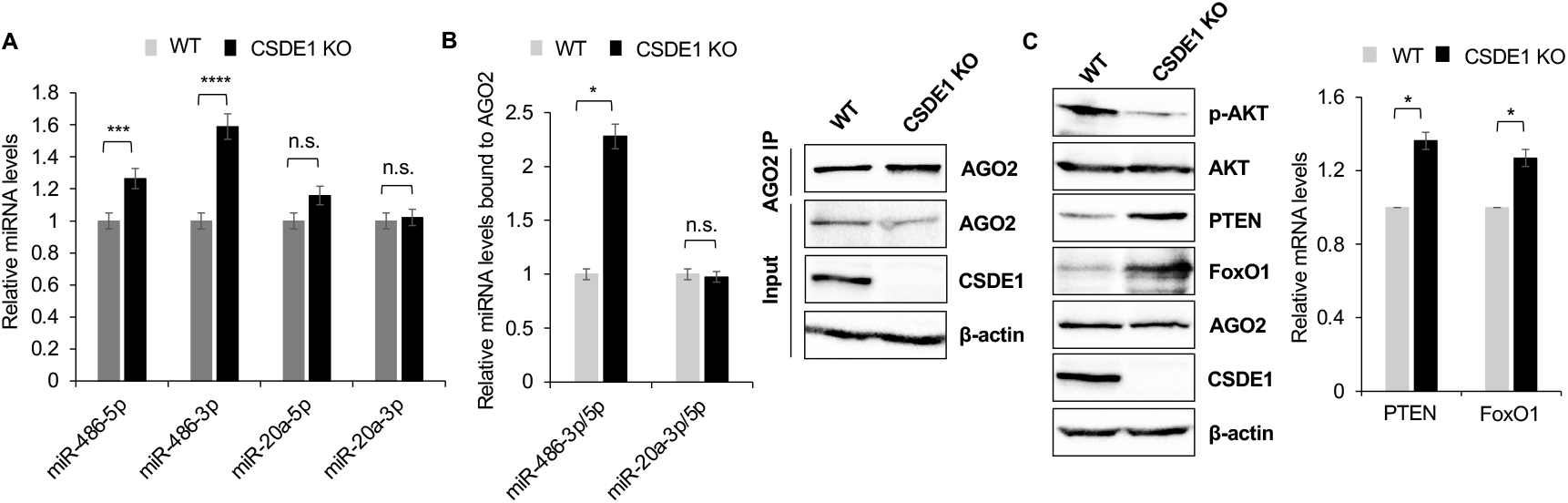
CSDE1 promotes miR-486-5p function in erythroleukemia cells. (A -B) Relative levels of miR-486-5p, miR-486-3p and canonical controls (miR-20a-5p and miR-20a-3p) in total RNA (A) and AGO2-bound (B) in the WT and CSDE1 KO MEL cells. (C) Western blot analysis of indicated proteins in WT and CSDE1 KO MEL cells (left), with relative mRNA levels of PTEN and FoxO1 (right). β-actin served as a housekeeping gene for relative mRNA quantification by dye-based RT-qPCR, and as a loading control for western blotting. U6snRNA served as a miRNA housekeeping gene for TaqMan miRNA assays. (*, p<0.05; **, p<0.01, ***, p<0.001, n=3, two tailed t-test; n.s., not significant, p>0.05).

We next examined the effect of CSDE1 on erythroid differentiation and apoptosis. Flow cytometry analysis comparing empty vector expressing cells with CSDE1 overexpressing cells revealed that CSDE1 expression reduced the median fluorescence intensity (MFI) of CD71, a marker of immature erythroid cells, while increasing the MFI of GlyA, a marker of mature erythroid cells (Figure S3A–B). These findings indicate that CSDE1 enhances erythroid maturation. Furthermore, CSDE1-expressing cells showed a higher proportion of Annexin V–positive cells (early apoptosis) and a lower proportion of 7-AAD–positive cells (late apoptosis), suggesting that CSDE1 facilitates the initiation of apoptosis while preventing progression to late-stage cell death. Collectively, these results demonstrate that CSDE1 promotes AGO2-mediated miR-486-5p expression and function in leukemia, while also supporting erythroid maturation and apoptosis regulation.

### The N-terminal domain of AGO2 regulates miR-486 strand ratio

Given that CSDE1 promotes miR-486-5p expression, we next asked whether CSDE1 influences strand selection through interaction with specific AGO2 domains. The N-terminal domain (N-domain) of AGO2 has been shown to facilitate release of the passenger strand (miR-486-3p) from the duplex, allowing the guide strand (miR-486-5p) to form mature RISC complexes (Kwak and Tomari 2012). To investigate this, we utilized wild-type (WT) AGO2 along with domain-deletion mutants (ΔN, ΔMID-PIWI, ΔPIWI) (Figure 2A). Western blot analysis of HA-immunoprecipitated samples confirmed that CSDE1 interacts with the AGO2 N-domain (Figure 2B). Notably, cells expressing the N-domain deletion mutant (ΔN) exhibited a significantly higher 3p/5p ratio in both total RNA and HA-IP fractions compared to WT AGO2 (Figure 2C–D). These results demonstrate that the AGO2 N-terminal domain plays a central role in regulating the miR-486-3p/5p ratio, thereby influencing strand selection.

**Figure 2.**
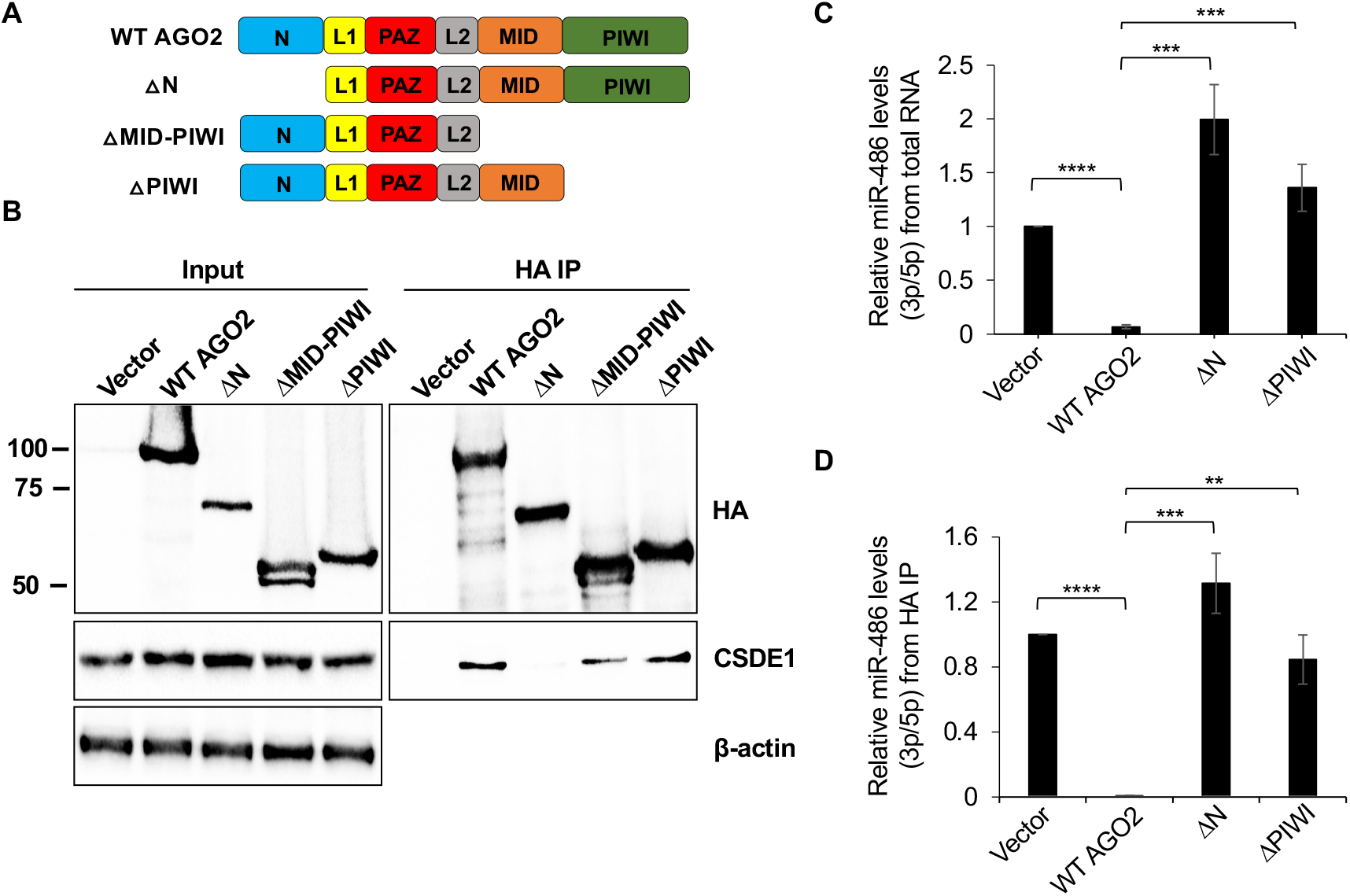
The AGO2 N-terminal domain interacts with CSDE1 and regulates the miR-486 strand ratio. (A) Schematics of AGO2 domains and its deletion mutants (ΔN, ΔMID-PIWI and ΔPIWI). (B) Western blots of the indicated proteins, from the IP of HA-tagged WT AGO2 and its mutants. (C-D) Relative miRNA levels in total RNA (C) and AGO2 IP (D) in HEK293T cells expressing WT AGO2 or its deletion mutants. U6snRNA served as the housekeeping gene for relative miRNA quantification, while β-actin served as a loading control for input in western blotting (**, p<0.03; **, p<0.01, ***, p<0.001, n=3, two tailed t-test).

### CSDE1 promotes miR-486 passenger strand cleavage

Since CSDE1 interacts with the AGO2 N-terminal domain (Kakumani et al. 2020) (Figure 2B), we hypothesized that it may facilitate the catalytic removal of the miR-486-3p passenger strand from the miR-486 duplex. To test this, we performed in vitro cleavage assays using synthetic miR-486 duplexes in which the passenger strand (miR-486-3p) was radiolabeled with ^32^P at its 5′ end (Figure 3A). The reaction mixtures were resolved on Urea-PAGE gels to visualize cleavage products. Cleavage assays and time-course experiments using CSDE1 KO lysates revealed a marked reduction in miR-486-3p passenger-strand cleavage efficiency, with the cleavage ratio (band intensity of the cleaved fragment relative to the intact 3p strand) being significantly higher in WT compared to CSDE1 KO samples (Figure 3B–C). Reintroduction of full-length CSDE1 into KO cells restored cleavage activity to near-wild-type levels, whereas KO lysates transfected with CSDE1 KO (Mock) remained defective (Figure 3D). Consistently, time-course analysis confirmed progressive recovery of cleavage activity in the CSDE1 rescue condition (Figure 3E). These findings demonstrate that CSDE1 enhances AGO2-mediated slicer-dependent removal of the miR-486 passenger strand.

**Figure 3.**
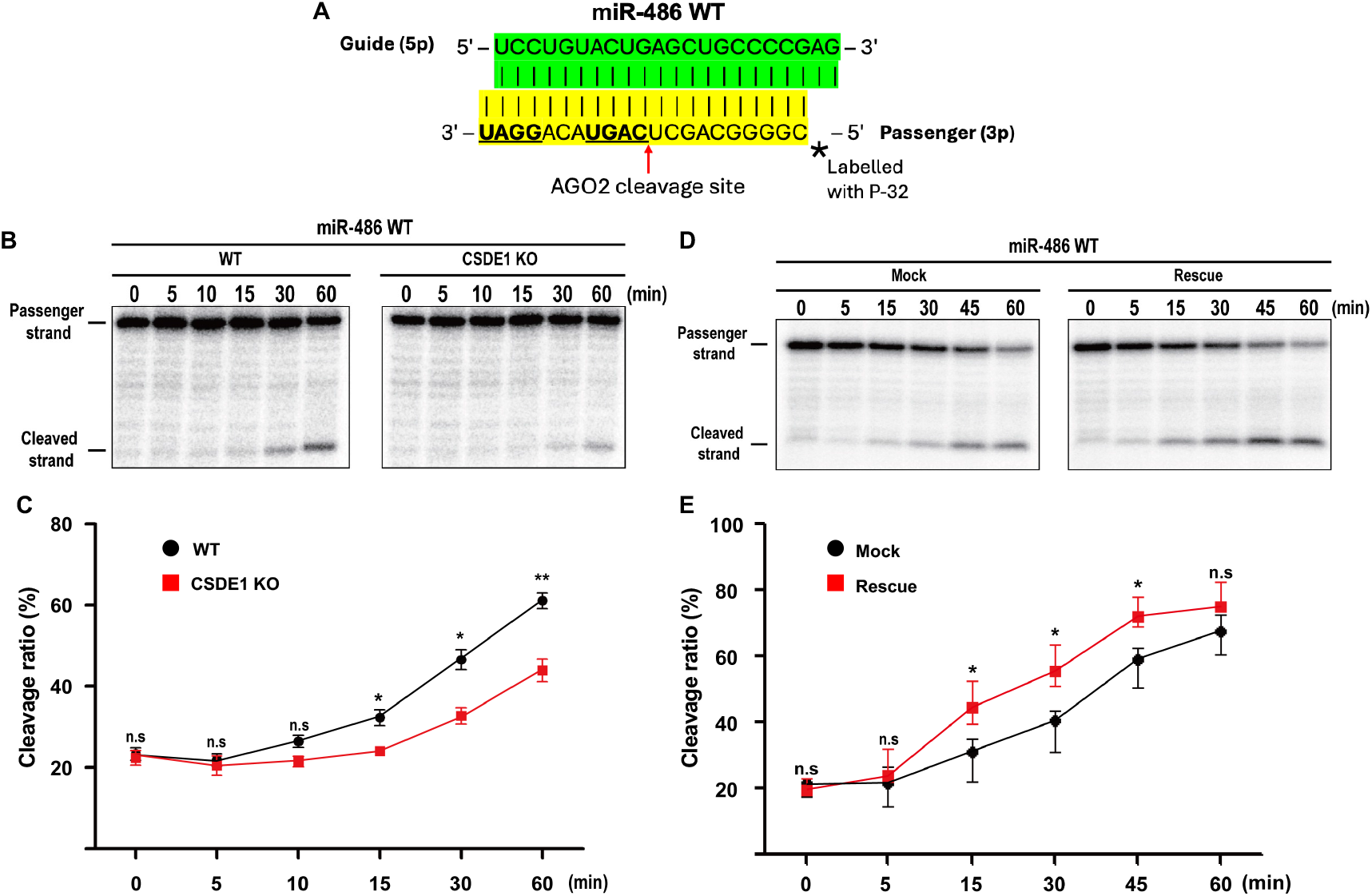
CSDE1 promotes AGO2-dependent cleavage of the miR-486-3p passenger strand. (A) Schematic representation of synthetic miR-486 duplexes used in the in vitro cleavage assay, in which the passenger strand (miR-486-3p) was radiolabelled with P-32 at the 5′ end. (B) A representative autoradiograph of Urea-PAGE from the in vitro cleavage assay showing cleaved products in the WT and CSDE1 KO cell lysate at 20°C. (C) Line graph of cleavage ratios (intensity of the cleaved band relative to the intact strand) in the WT and CSDE1 KO cell lysate. (D) A representative autoradiograph showing cleavage efficiency in the lysate of CSDE1 KO cells transfected with full-length CSDE1 (Rescue) or vector control (Mock). (E) Line graph of cleavage ratios from the time-course in vitro cleavage assay at 25°C (*, p<0.05; **, p<0.03; n=3; n.s., not significant, p>0.05).

Furthermore, to determine whether CSDE1 exhibits strand selectivity, we examined the miR-486 sequence and identified a conserved CSD motif (5′-GGAU-3′) in the passenger strand (miR-486-3p) (Figure S4A). Analysis of relative miRNA expression data to the total RNA showed that CSDE1 preferentially associates with the miR-486-3p strand compared with the guide strand, miR-486-5p (Figure S4B). Importantly, the expression level of miR-486-3p was similar to that of miR-451, which is known to contain a CSD-binding motif recognized by CSDE1(Kakumani et al. 2023), thus indicating potential mechanistic basis for strand preference. Now, to assess the role of the CSD motif in CSDE1-mediated strand recognition, we generated two mutant versions of the miR-486-3p passenger strand (Mut1 and Mut2) in which the motif sequence was altered while preserving duplex integrity (Figure S5A). In vitro cleavage assays using these mutant duplexes revealed that the intensity of the band corresponding to cleaved miR-486-3p was markedly higher in the CSDE1 Rescue compared with the CSDE1 knockout (Mock) (Figure S5B). This was corroborated by time-course analyses, which showed a significantly higher cleavage ratio in the Rescue than in the Mock (Figure S5C). Notably, when cleavage ratios were compared between wild-type miR-486 and the Mut1/Mut2 variants, no significant differences were observed (Figure S5D). Together, these findings suggest that although CSDE1 selectively binds the miR-486-3p passenger strand and recognizes the CSD motif, this sequence alone is not essential for AGO2-mediated passenger-strand cleavage.

### The N-terminal CSD1 domain is crucial for miR-486 passenger strand cleavage

Previous studies have demonstrated that the N-terminal cold-shock domain 1 (CSD1) of CSDE1 is essential for its interaction with AGO2 and for its RNA-binding capacity (Kakumani et al., 2021; Hollmann et al., 2020). To directly assess the role of this domain in passenger-strand removal, we performed rescue experiments in CSDE1 knockout (KO) cells expressing either full-length CSDE1 or a ΔCSD1 mutant lacking the N-terminal domain. In vitro cleavage assays demonstrated that full-length CSDE1 restored miR-486 passenger-strand cleavage to wild-type levels, whereas the ΔCSD1 mutant failed to rescue activity and showed cleavage efficiency indistinguishable from the KO mock control (Figure 4A–B). Statistical analysis confirmed that the CSDE1 rescue condition was significantly different from both ΔCSD1 and mock, while ΔCSD1 and mock did not differ from each other (Figure 4C). These results demonstrate that the CSD1 domain is indispensable for CSDE1-mediated enhancement of AGO2-dependent passenger-strand cleavage, likely by enabling stable RNA binding and/or direct interaction with the AGO2 N-terminal domain to facilitate efficient catalytic strand separation.

**Figure 4.**
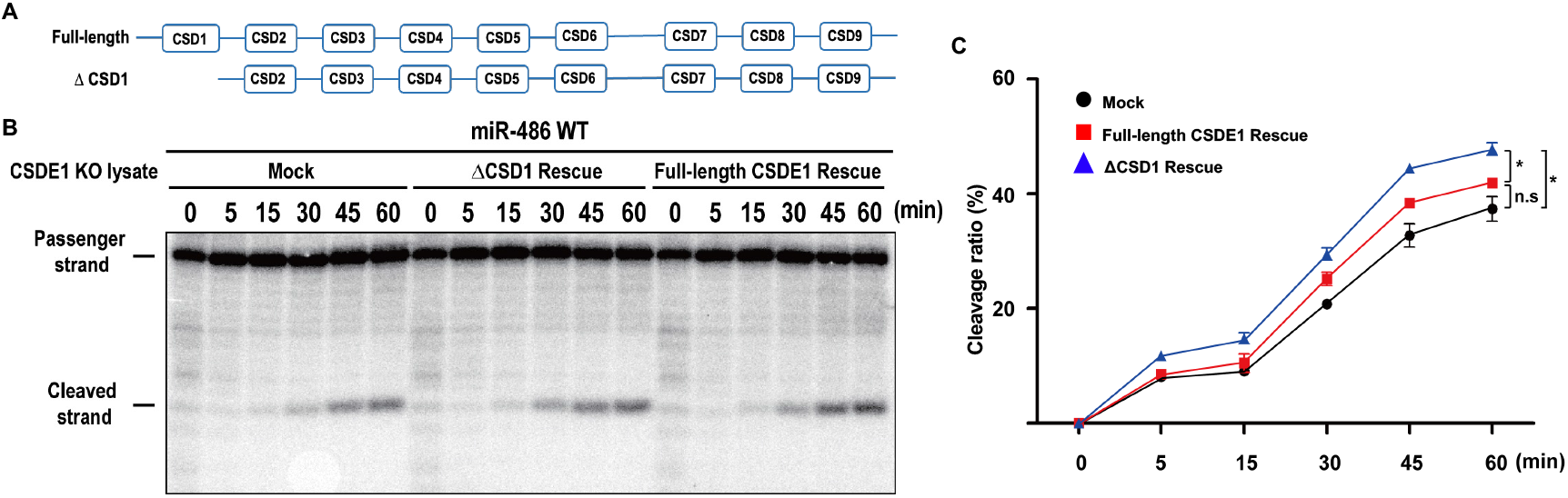
The N-terminal CSD1 of CSDE1 is essential for miR-486 passenger-strand cleavage. (A) Schematic representation of CSDs in CSDE1 and the N-terminal deletion mutant used in the study. (B) A representative autoradiograph of Urea-PAGE gel from in vitro cleavage assay performed in CSDE1 KO cells transfected with full length CSDE1 or ΔCSD1. (C) Line graph of cleavage ratios from the cleavage assay. (*, p<0.05; n=3, n.s., not significant, p>0.05).

## DISCUSSION

miRNA maturation is tightly regulated at multiple steps, and the balance between guide and passenger strands is a key determinant of miRNA function. While canonical miRNAs depend on Drosha and Dicer for duplex processing, a subset of non-canonical miRNAs including miR-486 relies on AGO2 slicer activity for passenger-strand removal (Kakumani et al. 2023; Jee et al. 2018). This process can be further influenced by RNA-binding proteins (RBPs) that stabilize precursors or modulate their engagement with AGO2 (Kakumani et al. 2023). In this study, we identify CSDE1 as a critical regulator of AGO2-mediated strand separation and maturation of miR-486-5p in leukemia. Our results demonstrate that CSDE1 is highly expressed in normal hematopoietic cells but downregulated in AML (Figure S1–S2). Loss of CSDE1 led to retention of the passenger strand (miR-486-3p), an elevated 3p/5p ratio, and derepression of FOXO1 and PTEN (Figure 1A–C), consistent with impaired miR-486-5p activity. Overexpression of CSDE1 promoted erythroid maturation, as reflected by reduced CD71 and increased GlyA expression, and facilitated early apoptosis while preventing late-stage cell death. These cellular outcomes align with the tumor-suppressive function of miR-486-5p and suggest that CSDE1 contributes to erythroid differentiation programs by reinforcing miR-486-5p activity. Functionally, this disruption impaired erythroid differentiation and apoptosis (Figure S3A–B), thereby promoting leukemic progression. These findings are summarized in our schematic model (Figure 6), which depicts how CSDE1 supports normal erythropoiesis, while its loss disrupts strand balance and promotes leukemic progression

Mechanistically, we provide evidence that CSDE1 interacts with the AGO2 N-terminal domain (Figure 2A), which is known to be essential for duplex unwinding (Kwak and Tomari 2012). Notably, the ΔN-terminal AGO2 mutant disrupted CSDE1 binding and phenocopied CSDE1 loss, yielding elevated 3p/5p ratios and reduced guide-strand activity (Fig. 2B–D). These findings suggest that the AGO2 N-domain provides a critical interface for CSDE1 to stabilize strand separation. Furthermore, because CSD1 is required for RNA binding by CSDE1 (Kakumani et al. 2020; Brandmann et al. 2018), we had hypothesized that CSDE1 simultaneously engages the passenger strand through CSD1 and AGO2 through its N-domain interface. We further investigated the role of the RNA-binding CSD1 domain of CSDE1. Rescue experiments revealed that while full-length CSDE1 significantly restored miR-486 cleavage efficiency, the ΔCSD1 mutant failed to do so, showing no difference from CSDE1 knockout (Figure 4A-C). These results demonstrate that CSD1 is essential for efficient passenger-strand removal. Taken together, our data suggest a model in which CSDE1 engages AGO2 through its N-terminal interface while simultaneously binding RNA via CSD1. This dual interaction may couple strand recognition with AGO2 catalytic cleavage, thereby accelerating passenger-strand clearance and reinforcing guide-strand fidelity. Collectively, our findings establish CSDE1 as a key enhancer of AGO2-dependent miR-486-5p biogenesis in erythroleukemia. By binding both AGO2 and the passenger strand, CSDE1 ensures efficient guide-strand maturation, thereby linking RNA processing to lineage-specific differentiation and apoptosis. These insights expand the repertoire of non-canonical miRNA regulators and provide a mechanistic framework for understanding how post-transcriptional control influences hematopoietic fate.

From a translational perspective, our findings intersect with evolving therapeutic strategies in AML. Current treatment paradigms are shifting beyond intensive chemotherapy toward targeted approaches, including FLT3, IDH1/2, and BCL2 inhibitors, as well as venetoclax-based combinations, immune checkpoint inhibitors, and epigenetic therapies (Liu 2021; Wysota et al. 2024). However, drug resistance, clonal heterogeneity, and impaired differentiation remain major obstacles (Wysota et al. 2024). Our data suggest that CSDE1 loss undermines the tumor-suppressive activity of miR-486-5p, thereby sustaining leukemic self-renewal. Restoring CSDE1 function or mimicking its strand-separation activity may complement existing strategies by reactivating miRNA-mediated repression of oncogenic pathways. Pharmacologic agents or RNA mimetics designed to stabilize CSDE1–AGO2 interactions could, in principle, enhance RISC assembly and restore apoptotic and differentiation programs downstream of miR-486-5p. Future investigations must define the broader clinical significance of CSDE1 in hematological malignancies. Its reduced expression in AML raises the possibility that genetic or epigenetic lesions in CSDE1 contribute to leukemogenesis or therapy resistance, with mutational analysis of domains such as CSD1 offering a path to identify variants that disrupt AGO2 engagement and strand clearance. At the mechanistic level, key questions remain regarding the structural features that confer CSDE1 selectivity for AGO2-dependent duplexes, and whether additional cofactors such as helicases or exonucleases cooperate in passenger-strand removal. Importantly, in vivo studies are needed to test whether CSDE1 deficiency mirrors the phenotypes of AGO2 catalytic mutants or miR-451/486 double knockouts, thereby clarifying its role in maintaining hematopoietic homeostasis.

In conclusion, our study establishes CSDE1 as an essential regulator of AGO2-mediated miR-486 maturation. By coupling strand separation to erythroid differentiation and tumor suppression, CSDE1 integrates RNA processing with hematopoietic fate decisions. These findings position the CSDE1–AGO2–miR-486 axis as a promising target within the rapidly expanding therapeutic space of AML, with broad implications for the post-transcriptional control of cancer biology.

## MATERIALS AND METHODS

### Cell culture and transfection

HEK293T and MEL cells were grown in Dulbecco’s modified Eagle’s medium (DMEM) and Roswell Park Memorial Institute (RPMI) 1640 Medium respectively supplemented with 10% fetal bovine serum, 50 U/ml penicillin, and 50 μg/ml streptomycin, respectively. K562 cells were cultured at a density of 0.5M-1M cells/mL in RPMI 1640 with L-Glutamine (Wisent Bioproducts, 350-000-CL) supplemented with 10% FBS (Wisent Bioproducts, 098-150). LentiX cells were cultured in DMEM (Wisent Bioproducts, 319-015-CL) supplemented with 10% FBS and 1mM sodium pyruvate (Wisent Bioproducts, 600-110-EL). The transfections in HEK293T cells were performed using jetPRIME and jetOPTIMUS reagents following the manufacturer’s instructions (Polyplus transfection SA, Illkirch, France).

### Lentivirus preparation, transduction

Lentiviral particles were produced as previously described (REFERENCE: doi: 10.1158/2643-3230.BCD-22-0086). Briefly, LentiX cells were transfected with packaging plasmids (psPAX2, pMD2.G) and lentiviral overexpression vectors using Lipofectamine 2000 (ThermoFisher, 11668500). Viral supernatant was collected after 48 hours, filtered through 0.45 um filters, concentrated by ultracentrifugation, and snap frozen. Lentiviruses were titrated on HeLa cells. K562 cells were infected by adding concentrated lentivirus directly to the culture media at a multiplicity of infection (MOI) of 2-3 infectious units/cell.

### Flow cytometry analysis

For surface immunophenotyping of lentivirally infected K562 cells, cells were washed in FACS buffer (PBS + 2% FBS + 2mM EDTA) and stained with a CD71-UV496 (BD Biosciences, 750652, used at 1:100) and GlyA-SB780 (Thermo Fisher, 78-9987-42, used at 1:100) for 20 minutes at room temperature. Cells were then washed and resuspended in FACS buffer with 1 ug/mL DAPI for flow cytometry. For apoptosis analysis of K562 of lentivirally infected K562 cells, cells were washed in 1x AnnexinV Binding Buffer (prepared from 10x concentrate, BD Biosciences, 556454) and stained with AnnexinV-AF647 (ThermoFisher, A23204, used at 1:75) and 7AAD (BD Biosciences, 559925, used at 1:75 for 20 minutes at room temperature. Cells were washed and resuspended in 1x AnnexinV Binding Buffer with 1:75 7AAD for flow cytometry. Flow cytometry was performed on a BD LSRFortessa™ Cell Analyzer. Data were analyzed using FlowJo (v10.10.0).

### Co-immunoprecipitation

Cells were homogenized in the lysis buffer with protease and phosphatase inhibitor cocktail. The lysate was centrifuged at 15 000g for 15 min, the supernatant was collected and measured for protein concentration using Bradford reagent. Meanwhile, Dynabeads Protein G (Thermofisher) were prepared in an Eppendorf (25 μl for a total protein extract of 2 mg and 10 μg of antibody). The beads were washed with 3 times the volume of lysis buffer and was repeated two more times. The lysate was preincubated with the equivalent of Dynabeads at 4°C on a nutator for about 45 min. The lysate was collected, and the respective antibody added to incubate on nutator at 4°C for overnight. The next day, Dynabeads were prepared (as stated previously), and the lysate was added to incubate on the rotator at room temperature for about 90 min. For RNase treatment, the RNases A&T1 were added at a concentration of 4 μg and 1 unit respectively per 1 mg of total protein extract in buffered solution to the beads and incubated at RT for 15 min. The beads were washed about 5 times with lysis buffer, and the samples were extracted for western blotting by adding SDS loading buffer in 1× concentration and heating at 100°C. For Northern blotting, the samples were extracted using Trizol as per manufacturer’s instructions (Thermofisher).

### Quantitative RT-PCR

Total RNA extraction was performed using TRI Reagent^®^ as per the manufacturer’s instructions (Sigma-Aldrich). mRNA expression analysis was performed using the GoTaq® 1-Step RT-qPCR System following the manufacturer’s protocol (Promega). Each reaction contained 200 ng of RNA template. Primers were designed through Integrated DNA Technologies, and β-actin was used as the internal reference control. To measure miRNA levels, reverse transcription using total RNA, followed by quantitative PCR was performed using the TaqMan™ Universal PCR Master Mix (ThermoFisher Scientific) in accordance with the manufacturer’s guidelines. Assays were conducted for miR-486-5p, miR-486-3p, miR-20a-5p, miR-20a-3p and miR-451 with U6 snRNA serving as the endogenous control. In both the cases, relative expression levels were measured using the ΔΔCt method and analyzed with BioRad CFX Maestro Software and Microsoft Excel.

### *In vitro* passenger-strand cleavage assay

*In vitro* passenger-strand cleavage assays were performed using 293T WT cell lysate expressing hAOG2, and CSDE1 KO cell lysates co-expressing hAOG2 with MOCK, ΔCSD1, or CSDE1. *In vitro* passenger-strand cleavage assays were performed by previously described method with some modifications (Park and Shin 2015). Briefly, 3μL cell lysate, 1.5μL reaction buffer, and 0.5μL of 100 nM 5′-^32^P-radiolabeled miR-486-3p-WT or miR-486-3p-CSDM were used in the assay. The reaction samples were incubated at 25°C for indicated time. To stop the reaction, samples were snap-frozen in liquid nitrogen, then mixed with 7.5μL of 2X formamide loading dye and heated at 95°C for 5 minutes. The resulting mixtures were analyzed on a 15% denaturing polyacrylamide gel. Cleaved passenger strands were detected by Typhoon FLA-7000 image analyzer (Fujifilm Life Sciences) and quantified with ImageJ.

## ACKNOWLEDGEMENTS

We thank all members of our laboratories for their input helpful discussions and grateful to Prof. Marieke von Lindern, Sanquin Research, Amsterdam for the generous gift of MEL WT and CSDE1 KO cell lines.

## FIGURE LEGENDS

**Figure S1.**
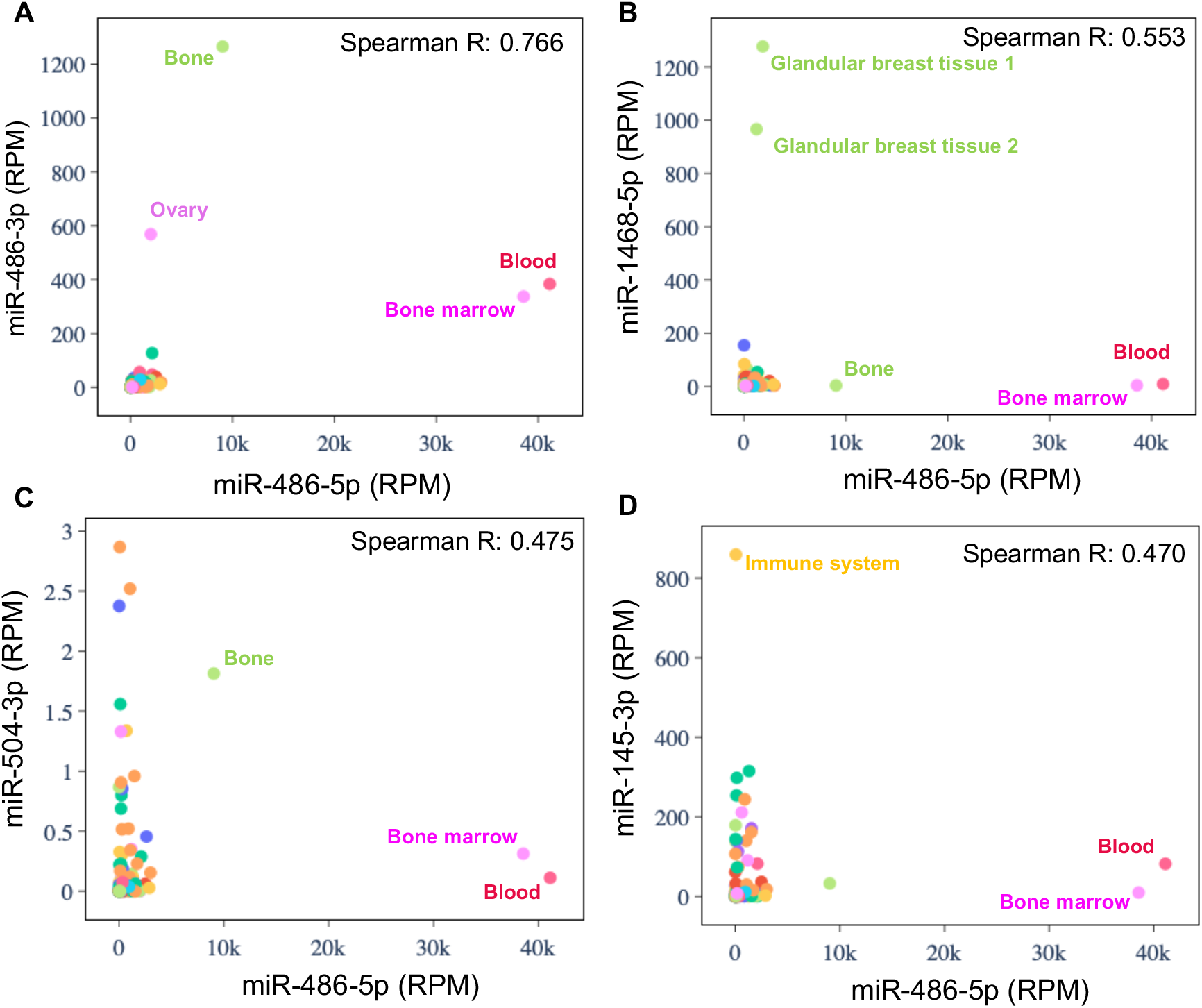
miR-486-5p is highly expressed in blood, relative to miR-486-3p and the other canonical miRNAs. (A-D) Scatterplots show correlations between the expression levels of miR-486-5p vs miR-486-3p (A), or other canonical miRNAs, namely miR-1468-5p (B), miR-504-3p (C), and miR-145-3p (D) among different tissue types. The miRNA expression is shown in Reads Per Million (RPM), with the degree of correlation indicated by the Spearman coefficient (R).

**Figure S2.**
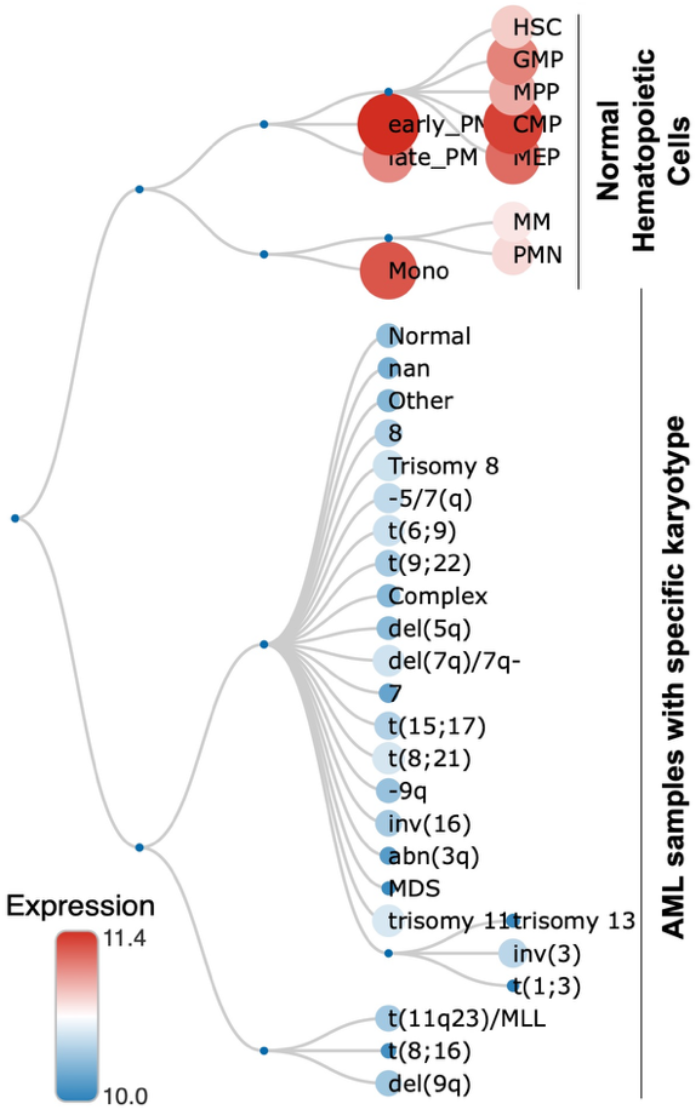
CSDE1 expression is downregulated in AML samples compared to normal hematopoietic cells. The hierarchical tree shows the relationship between the displayed samples, and the relative CSDE1 expression (log2 scale) is plotted based on curated microarray data.

**Figure S3.**
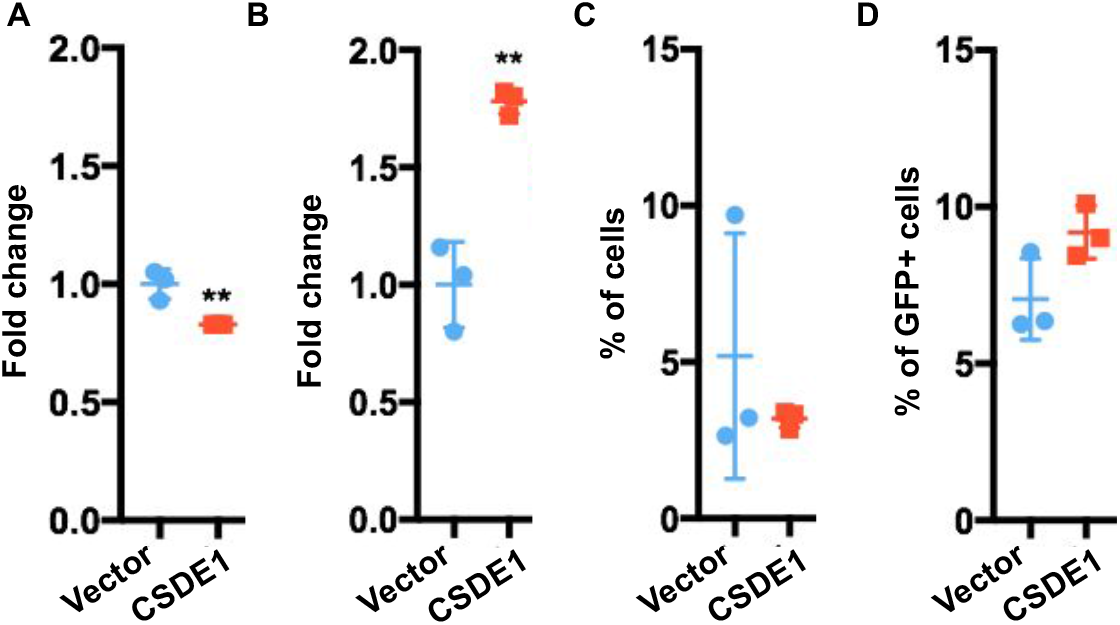
CSDE1 promotes erythroid maturation in erythroleukemia. (A-B) Flow cytometry analysis measuring the medium fluorescence intensity (MFI) of CD71 (A) and GlyA (B) in K562 cells transduced with lentiviruses expressing full-length CSDE1 (red) or the vector control (blue). (C-D) Apoptotic analysis under the same settings using cells labelled with 7-AAD (shown as a % of all cells) or Annexin V (shown as a % of GFP+ cells). (**, p<0.03; n=3, n.s., not significant, p>0.05).

**Figure S4.**
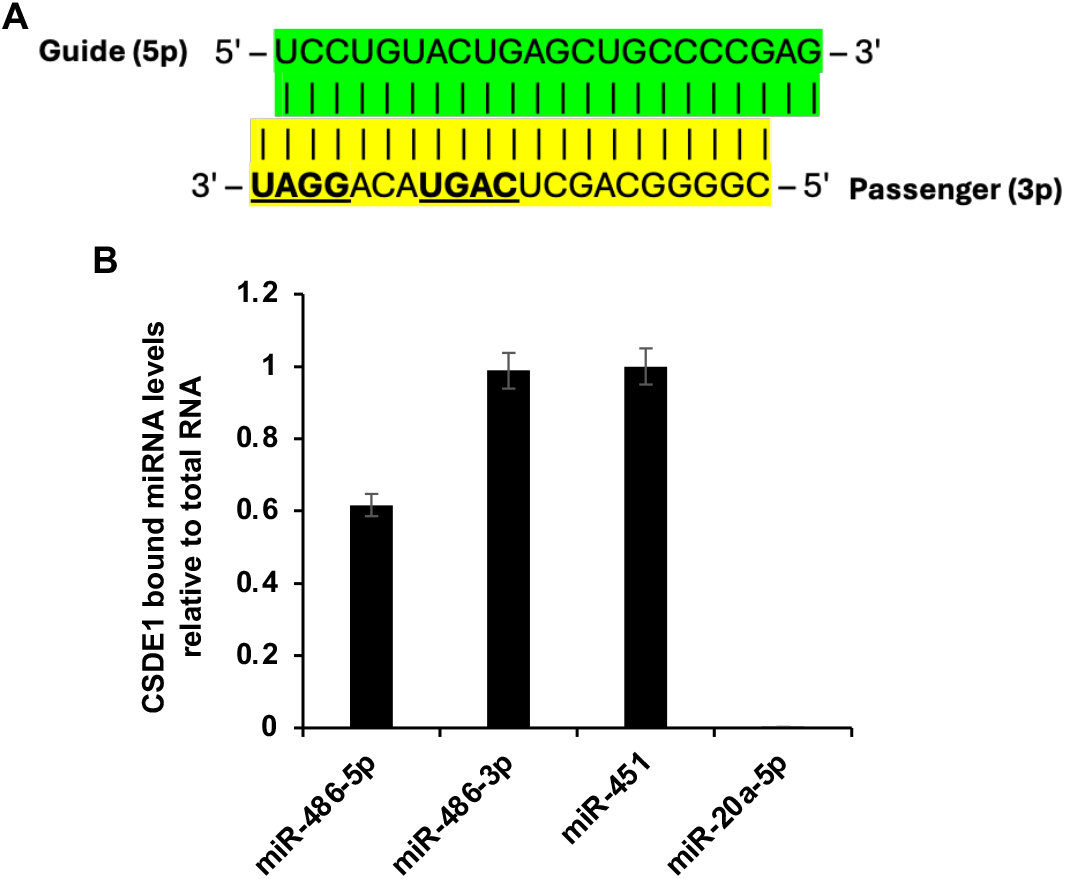
CSDE1 binds miR-486 duplex. (A) Sequence schematic highlighting potential CSD-binding motifs (5′-GGAU-3′, 5’-CAGU-3’) present in miR-486-3p. The CSD motif sequences are in bold and underlined. (B) Relative miRNA levels bound to CSDE1, compared to IgG control, and normalized to total RNA, in MEL cells.

**Figure S5.**
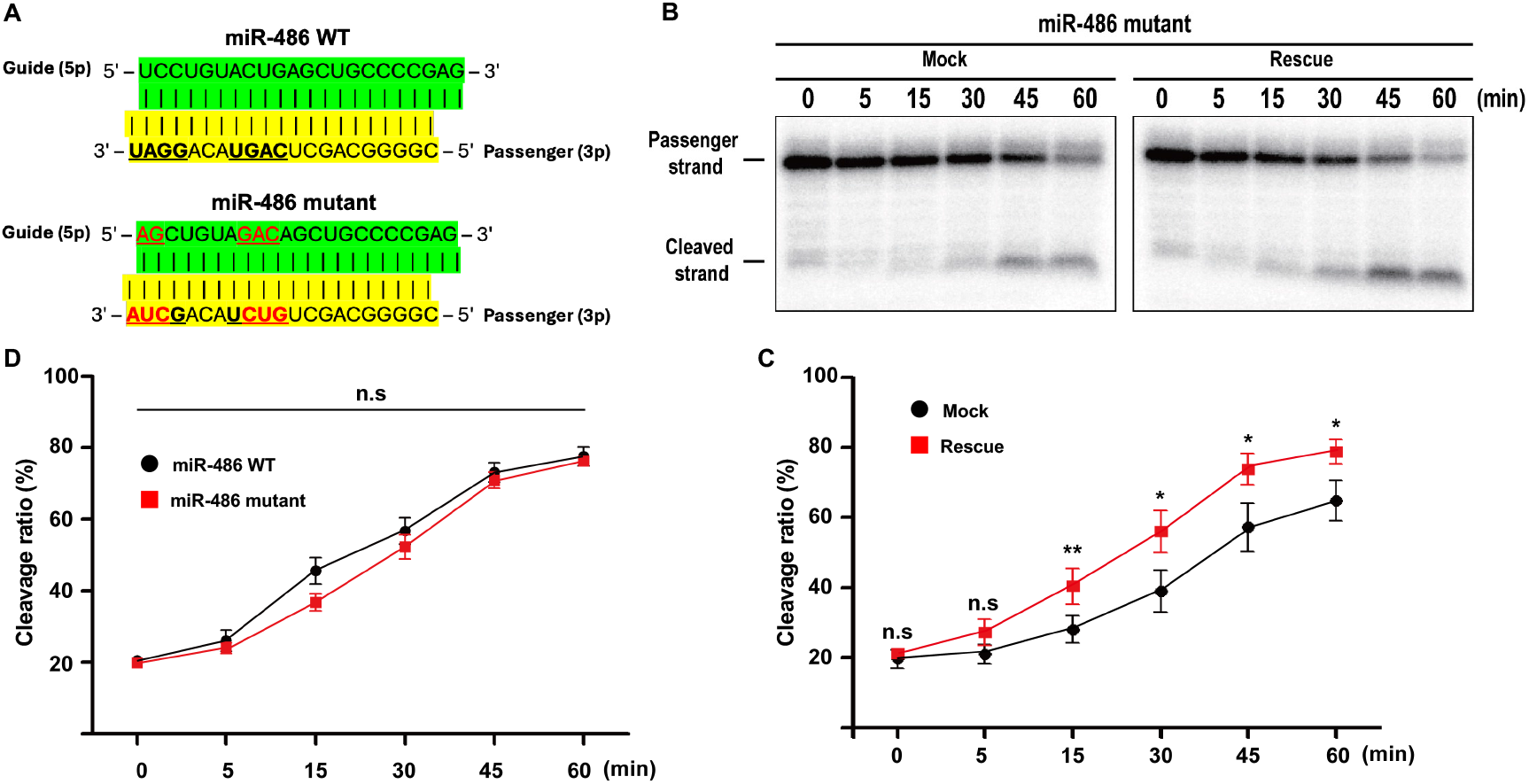
The CSD motif in miR-486-3p is dispensable for AGO2-mediated passenger-strand cleavage. (A) Schematics of the WT and mutant miR-486 duplexes used in the cleavage assay. The CSD motif sequences are in bold and underlined. Nucleotide sequence in the CSD motifs were altered (in red) in the 3p to disrupt the site, and to maintain full complementarity between 5p and 3p strand, corresponding nucleotides in the 5p were also altered (in red). (B) A representative autoradiograph of Urea-PAGE gel from the in vitro cleavage assay using the lysate of CSDE1 KO cells under Resue or Mock conditions. (C) Line graph of cleavage ratios under the aforementioned experimental settings. (D) Line graph showing the comparison between cleavage ratios from CSDE1 rescue conditions supplied with the WT and the CSD mutant miR-486 duplexes. (*, p<0.05; **, p<0.03; n=3, n.s., not significant, p>0.05)

